# Tandem WW/PPxY motif interactions in WWOX: the multifaceted role of the second WW domain

**DOI:** 10.1101/2021.12.01.470705

**Authors:** Shahar Rotem-Bamberger, Jamal Fahoum, Keren Keinan-Adamsky, Tomer Tsaban, Orly Avraham, Deborah E. Shalev, Jordan H. Chill, Ora Schueler-Furman

## Abstract

Class I WW domains mediate protein interactions by binding short linear PPxY motifs. They occur predominantly as tandem repeats, and their target proteins often contain multiple PPxY motifs, but the interplay of WW/peptide interactions is not always intuitive. WW domain-containing oxidoreductase (WWOX) protein harbors two WW domains: unstable WW1 capable of PPxY binding, and well-folded but mutated WW2 that cannot bind such motifs. WW2 is considered to act as a WW1 chaperone, but the underlying mechanism remains to be revealed. Here we combine NMR, ITC and structural modeling to elucidate the role of both WW domains in WWOX binding to single and double motif peptides derived from its substrate ErbB4. Using NMR we identified an interaction surface between the two domains that supports a WWOX conformation that is compatible with peptide substrate binding. ITC and NMR measurements reveal that while binding affinity to a single motif is marginally increased in the presence of WW2, affinity to a dual motif peptide increases tenfold, and that WW2 can directly bind double motif-peptides using its canonical binding site. Finally, differential binding of peptides in a mutagenesis study is consistent with a parallel orientation binding to the WW1-WW2 tandem domain, agreeing with structural models of the interaction. Our results reveal the complex nature of tandem WW domain organization and substrate binding, highlighting the contribution of WWOX WW2 to both stability and binding. This opens the way to assess how evolution can utilize the multivariate nature of binding to fine-tune interactions for specific biological functions.

## Introduction

WW domains are small 38-40 residue modules that adopt a three-stranded antiparallel β-sheet fold. They are named for their two conserved tryptophan (W) residues. The first W helps stabilize the domain, while the second W facilitates binding to short, linear motifs rich in prolines (P), (1–3). WW domains mediate protein-protein interactions involved in a range of protein functions, from degradation by ubiquitination to nuclear transport. These protein functions are instrumental in determining cell fate by mediating apoptosis and cell proliferation.

WW domains usually occur in tandem, allowing for fine-tuned regulation through a combination of binding events (4). This complexity is enhanced by the presence of multiple proline-rich binding motifs, such as the class I PPxY motif, on partner proteins. Many of the reported interactions between WW domain proteins and PPxY motif partners involve more than one WW domain-PPxY motif pair. For example, the tandem WW domain of Yki, the *Drosophila melanogaster* homolog of human YAP, simultaneously binds two Tgi PPxY peptide motifs. Similarly, the Nedd4 WW2WW3 and WW3WW4 tandem domains bind a double motif-containing peptide derived from ARRDC3 (5,6). Moreover, the WW tandem domains of proteins, such as formin-binding protein 21 (FBP21), YAP and TAZ, bind dual PPxY peptide motifs with higher affinity than a corresponding single PY peptide motif (7,8). Thus, these multiple interactions enhance the binding affinity compared to single peptide motif binding (5,9), but also allow for fine tuning of binding, as for example in the protein kidney and brain (KIBRA) (10). It was also shown that while a minimal stretch that contains the three PPxY motifs of NDFIP2 retains the ability to activate ITCH, mutation of any one of the PPxY elements reduces activity (11).

Human WW domain containing oxidoreductase (WWOX) is a tumor suppressor involved in many biological functions such as apoptosis, DNA damage, inhibition of cell growth, cellular metabolism and proper neurodevelopment (12–14). WWOX contains two N-terminal WW domains (WW1 & WW2) separated by a short linker, as well as a C-terminal short-chain dehydrogenase/reductase (SDR) domain (**Figure 1**) (15,16). WWOX binds many class I proline rich motif-containing proteins, including ErbB4 and p73 (17–20), and competition with other WW domain proteins such as YAP for these binding partners has been shown to influence cellular behavior. Thus, WWOX sequesters ErbB4 in the cytosol by preventing its binding to YAP and its subsequent shuttling to the nucleus for the initiation of a transcriptional program leading to cellular proliferation (20). While it would seem that the two WWOX domains are a prototypical case of the dual interactions described above, the second of these, WW2, is in fact considered incapable of independently binding PPxY peptides with measurable affinity, due to the W85Y mutation in the conserved binding-pocket (In fact, the reverse Y85W mutation re-establishes binding for this domain (21)). WWOX therefore seems to bind PPxY partners exclusively through WW1. What then is the role of WW2, and how is WWOX able to compete with WW proteins with two fully functioning WW domains, especially for partner proteins that contain multiple PPxY motifs? It has been suggested that WW2 functions as a chaperone that stabilizes WW1 to improve the latter’s binding affinity to its partners (22), but the details of such a possible mechanism remain to be elucidated.

**Figure 1:**
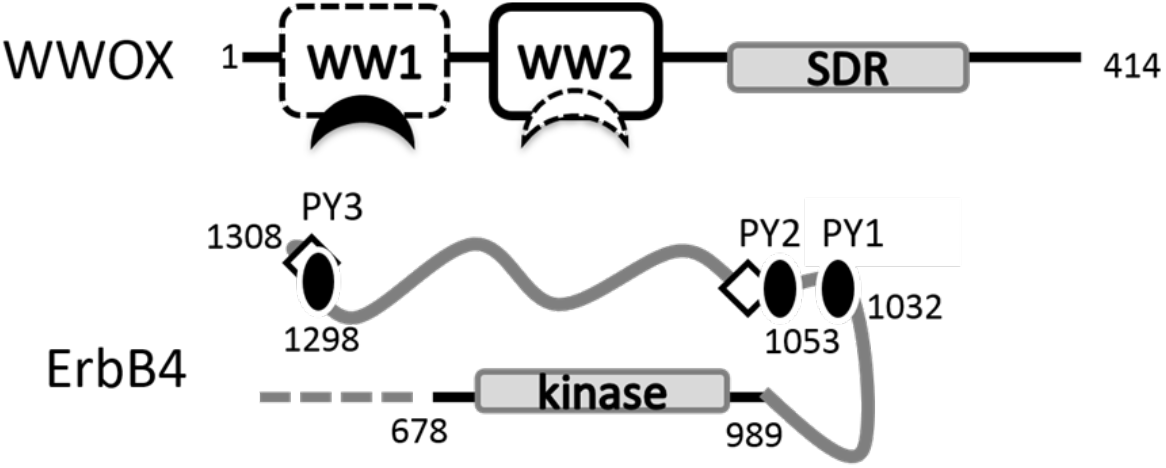
WWOX and its substrate ErbB4. WWOX contains two WW domains (residues 16-49 and 57-91, respectively), and an SDR domain (residues 122-383; domain position annotations according to InterPro (24)). Isolated WW1 is unstable (dashed lines) but binds to PPxY motifs (filled crescent), while WW2 is stable (solid lines), but was not reported to bind to PPxY motifs with measurable affinity (dashed crescent (25)). WWOX partner ErbB4 is cleaved upon maturation, releasing the C-terminal cytoplasmic region (residues 676-1308) that contains a kinase domain (positions 718-974) followed by an extended disordered region (gray curve) that contains several recognition motifs, including the three WWOX binding PPxY motifs investigated here (PY1 1032-1035, PY2 1053-1056 and PY3 1298-1301; black ovals), and several phosphorylated tyrosine residues (diamonds).

Here we have studied the structural and functional role of WWOX WW2 in binding WWOX partners, focusing on ErbB4 and its three PPxY motifs in the intracellular domain that is released to the cytoplasm by γ-secretase after receptor stimulation (23) **(Figure 1)**. Using CD and NMR experiments, we found that the presence of the WW2 domain, as well as the binding of the peptide, induce WW1 structural stabilization. Furthermore, our ITC and NMR protein-peptide binding assays show increased affinity of a double-PPxY peptide to the WWOX tandem WW domains when compared to a single PPxY peptide binding to the tandem domain, or to isolated WW1 binding a double-PPxY. This suggests that WW2 can engage a suitably oriented PPxY motif, despite its missing key tryptophan residue. Also, NMR spectra establish that the double PPxY peptide induces a significantly more structured conformation of WWOX than the single PPxY peptide, once again supporting direct involvement of WW2. By comparing affinities and chemical shift perturbation effects induced by native, engineered and mutated double-PPxY motif peptides we deduced the binding pose and directionality of two-site ligand binding and the contribution of different molecular determinants to affinity. In aggregate, this data suggests a plausible model of the relative orientation between the two WWOX WW domains, and reveals details of the interaction between WWOX WW2 and WW1 domains, and substrate single and double motif peptides, which are discussed in the functional context of WWOX-substrate interactions.

## Results

To elucidate the role played by WW2 in WWOX functionality, we investigated its influence on the stability and binding of WWOX. We compared single WW1 and WW2 domains to the tandem domain WW1-WW2 in terms of structural stability and binding affinity to different peptides derived from ErbB4, a known WWOX substrate (20) (see **Figure 1** and **Table I**).

**Table I:**
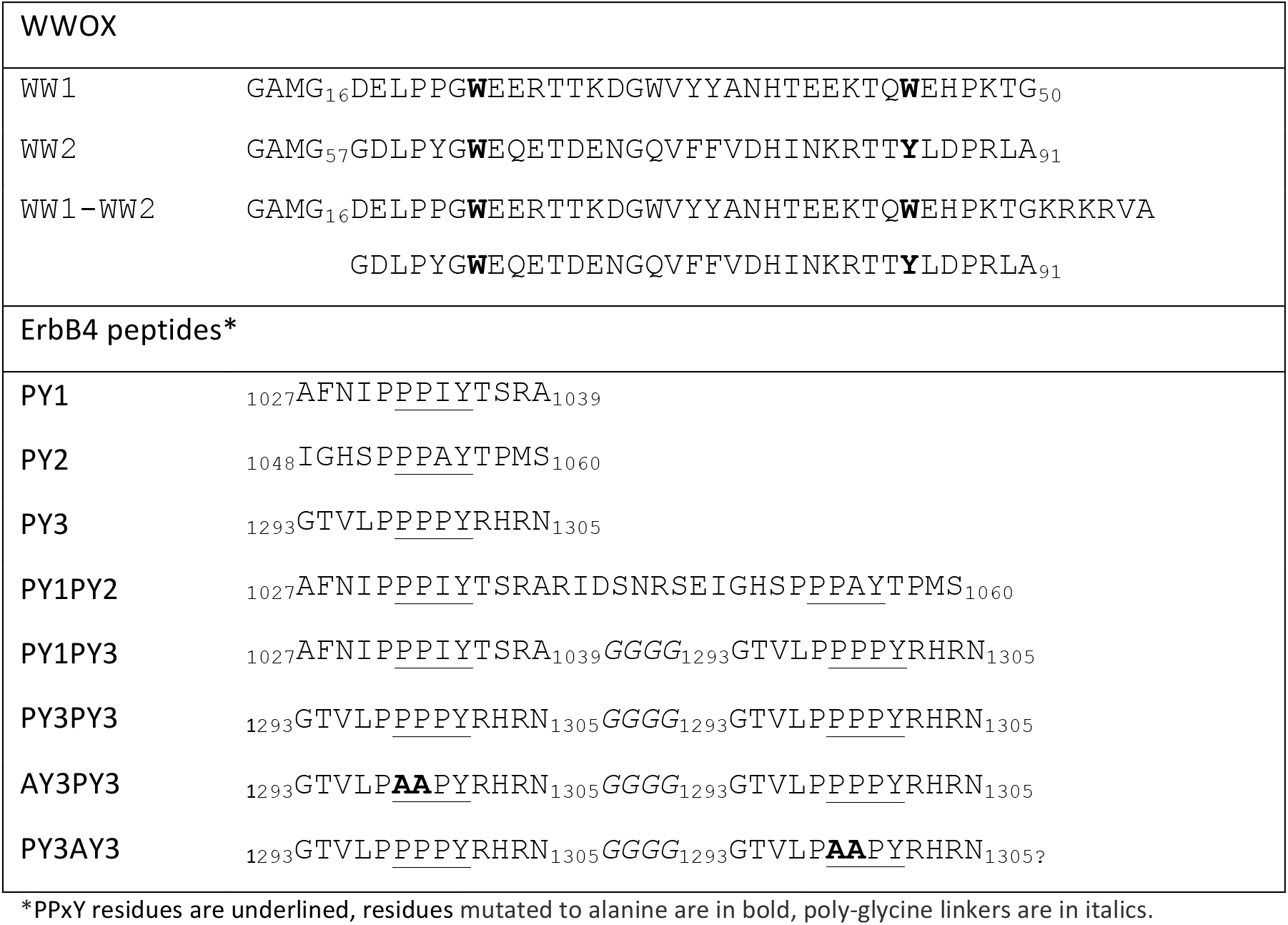
Proteins and peptides used in this study

### WWOX domain WW1 is stabilized by domain WW2

We used both denaturation experiments and CD spectra to characterize the stability of the isolated and tandem WWOX domains. Monitoring tryptophan fluorescence changes in urea denaturation experiments showed that WW1 is structured only within the context of the WW1-WW2 tandem domain, but not in its isolated form, indicating significant stabilization of WW1 by WW2 (**Figure 2A**). Folded WW domains exhibit a CD spectrum dominated by a strong positive peak at 225-230 nm contributed by ordered aromatic side chains, and a negative peak at >205 nm (26,27). The CD spectrum of isolated WW2 indeed conforms to this characteristic spectrum, albeit with a relatively weak positive peak and a negative peak at <205 nm, expected due to the W85Y mutation, while isolated WW1 is predominantly unstructured (**Figure 2B**). In comparison, two important spectral changes appear in the spectrum of tandem WW1-WW2: The WW1-WW2 positive peak is synergistically stronger than the sum of the WW1 and WW2 contributions, indicating mutual influence of the domains on each other resulting in a structure closer to a typical WW domain (26,27). Additionally, the negative band in the CD spectrum shifts to the right from 205 nm, indicating that the WW domain adopts a more folded, antiparallel β-sheet structure. Together these changes reinforce the view that WW2 significantly stabilizes WW1, as described before (22).

**Figure 2:**
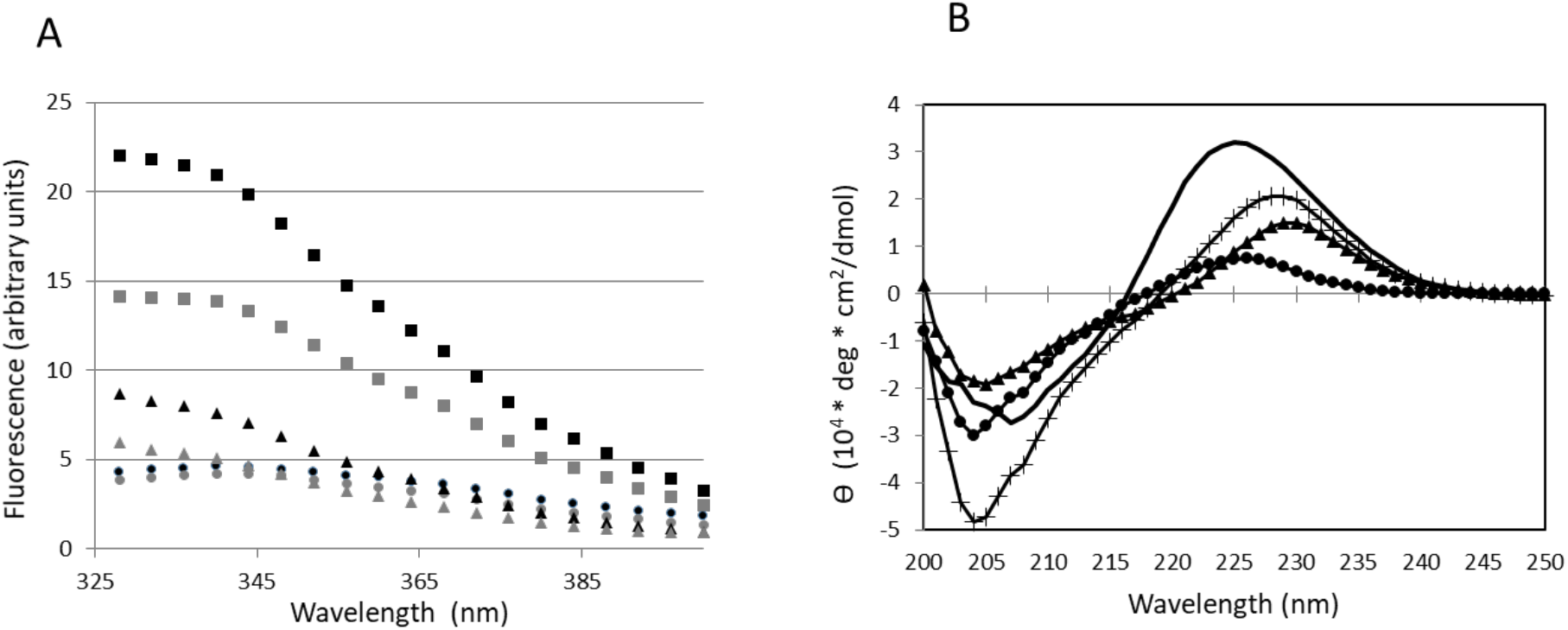
WWOX domain 1 (WW1) is stabilized by domain 2 (WW2). **A**. Tryptophan fluorescence assay for protein in native conditions (in black) and in 6 M urea (in grey). Whereas isolated WW1 (circles) exhibits little change, both WW2 and tandem WW1-WW2 domains (triangles and squares, respectively) exhibit significant change indicating a loss of structure. The larger change in the tandem domain as compared to WW2 indicates stabilization of WW1 by WW2. **B**. Circular dichroism curves for isolated WW1 (circles), isolated WW2 (triangles) and the tandem WW1-WW2 domains (line). Whereas WW1 is mostly unfolded, the latter two are ordered, showing a characteristic positive peak for WW domains around 225-230 nm. The tandem domain also shows a shift of the negative peak away from 205 nm, indicative of an extended β-sheet structure, and is significantly more structured than what would be observed from combining the spectra of the two individual domains (plus signs), highlighting the stabilization effect of WW2 on WW1.

### A molecular description of the WW2-WW1 interaction

For a more detailed picture of this stabilization event, we investigated its accompanying structural changes using well-established NMR-based methods. As ^1^H-^15^N HSQC spectra of proteins are highly sensitive to the local electronic environment, shifted cross-peaks in this ‘fingerprint’ spectrum (known as chemical shift perturbations, CSPs) report on local and global changes in structure and dynamics on a per-residue basis. The WW1-WW2 HSQC showed a striking presence of two sub-populations of cross-peaks, one exhibiting line-widths consistent with a small protein (below 10 kDa), and the other suffering from extensive line broadening that was aggravated at higher temperatures (data not shown). By recording spectra for the single domains we confirmed that the broadened peaks belong to WW1 residues while WW2 affords a well-behaved spectrum (**Figure 3A,B**). This broadening and its temperature-dependence suggested that WW1 loses signal intensity due to solvent-exchange, reflecting a less-folded and more flexible domain when compared to WW2.

**Figure 3:**
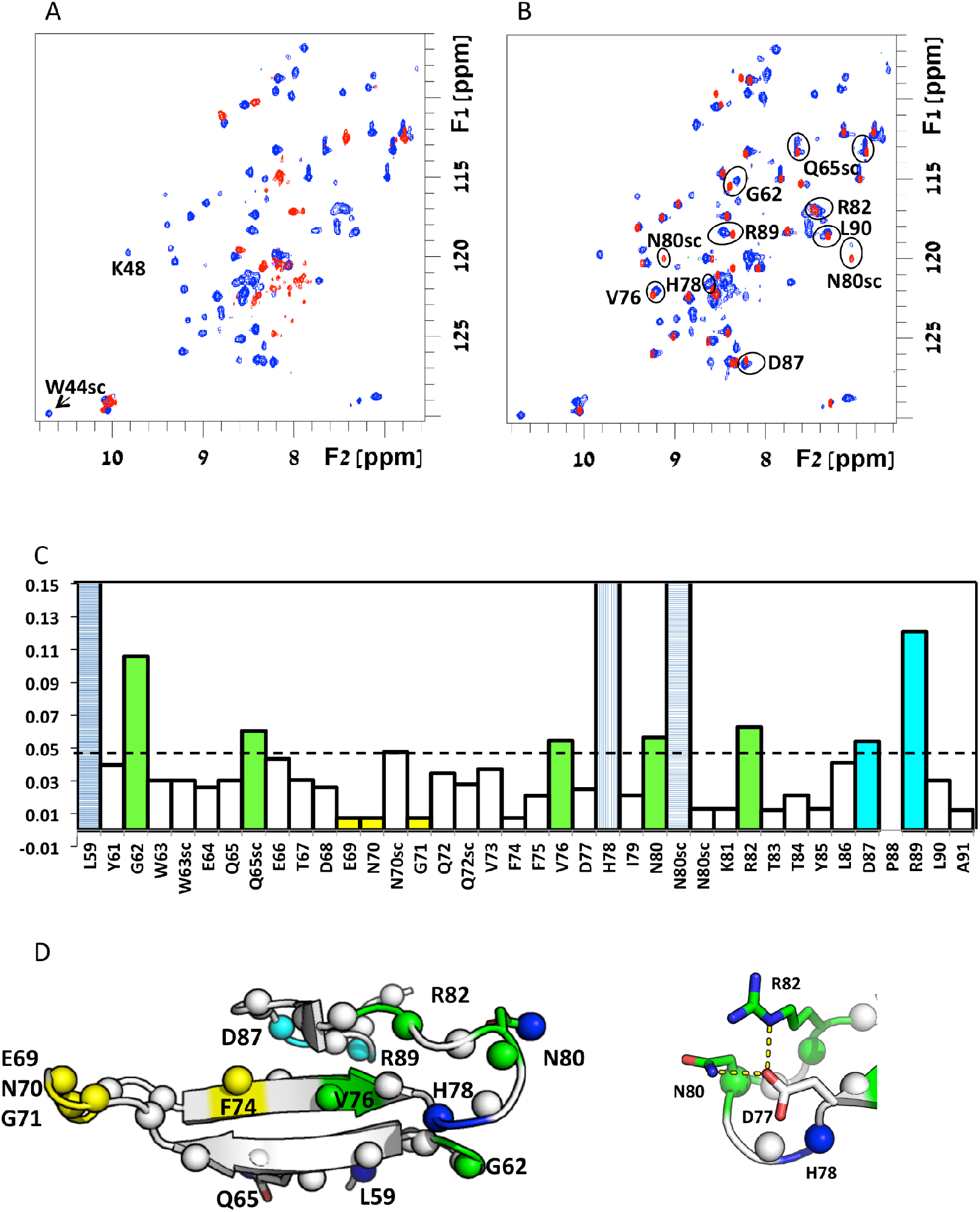
NMR HSQC spectra of single and tandem WWOX WW domains reveal an inter-domain stabilizing interaction. **A**. The ^1^H, ^15^N-HSQC spectrum of WW1 (red) shows a typical spectral dispersion of an unfolded domain, which changes to defined peaks in the WWOX tandem domain (blue), indicating that WW2 stabilizes WW1 (see **Supplementary Figure S1** for peak assignments). **B**. In contrast, structured WW2 (red) gives a well-behaved spectrum with many overlapping peaks with the WWOX tandem domain (blue). Peaks discussed in the text are annotated and/or highlighted by circles. **C**. Chemical shift perturbations (CSPs) of WW2 residues in the presence of WW1 (tandem domain). Significant CSPs are color-coded for their main regions, including the β2/β3 hairpin (green) and the C-terminal region (cyan); peaks that disappear/appear upon addition of WW1 (not fully assigned but clearly affected) are shown by blue (vertically/horizontally) hatched bars; stable peaks in the β1/β2 hairpin region are colored yellow. **D**. Mapping of changes onto the structure of the WW2 domain (PDB accession code 1WMV) using the same color scheme. Note the close vicinity of H78 (in blue) and G62 at the basis of the beta hairpin formed by strands β1 and β2 and other residues. Right, detailed structure of the β2/β3 hairpin. Figures of structures were generated using Pymol (28).

An analysis of differences between the two single-domain spectra and that of the WW1-WW2 tandem domain was instructive. WW1 cross-peaks in the tandem domain exhibited a significant global change, specifically an appearance of well-dispersed peaks and a reduction of peaks in the region characteristic for unstructured proteins, suggesting a stabilizing effect of WW2 upon the neighboring domain (**Figure 3A**). At the same time, WW2 peaks in two regions exhibited small yet significant intensity changes or cross-peak shifts (> 0.05 ppm, **Figures 3B,C**). One of these includes a cluster of residues on one side of the WW2 β-sheet, involving G62 (preceding β_1_) and the β_2_β_3_ turn (particularly H78, N80 and R82). Another region of interest is the second region consisting of C-terminal residues R89-L90-A91, whose chemical shifts suggest that this segment packs against the WW2 domain core, rather than adopting a free random-coil conformation, and that this interaction with the domain core is modified upon linking WW2 to WW1. These two effects could be attributed to direct interaction with WW1, or alternatively, to indirect effects propagated through the WW2 β-sheet. In summary, we find that the WW2 ‘tip’, comprised of pre-β_1_ and β_2_β_3_-turn residues plays an important role, or at least is significantly perturbed upon WW1 stabilization, and that the two WW-domain cores, linker and C-terminus form a contiguous set of interactions that further contribute to the WW1-WW2 structure.

### Binding of ErbB4 ligands to WWOX

How does the tandem domain bind its peptide substrates, and what does WW2 contribute to this affinity? Our previous study highlighted the variety of mutual influences between individual WW domains in tandem repeats upon substrate binding affinity and specificity (4). WWOX stood out due to the inability of its WW2 to bind to canonical substrates (e.g., PPxY motifs) at measurable affinities, due to a tryptophan-to-tyrosine mutation at the second characteristic W-position. To study the details of the suggested chaperone role of WW2 in WWOX binding of PPxY motifs we compared the affinities of isolated WW1 domain and tandem WW1-WW2 to various ErbB4-derived PPxY-containing peptides (see **Figure 1** and **Table I**). Affinities were measured using both ITC and NMR, a combination that allowed us to cover a wide range of affinities including often-inaccessible *K*_D_ values in the 200-1000 μM range.

In terms of affinity, ITC results differentiated between the affinity of tandem WW1WW2 to PY3 (*K*_D_ = 30 μM) and PY1/PY2 (estimated weak binding, *K*_D_ > 100 μM) (**Table II)**, in qualitative agreement with previous studies (22). However, although no significant affinity to the isolated WW2 domain was detected for either peptide (*K*_D_ >> 200 μM), ITC showed an approximate two-fold decrease in binding affinity when the WW2 domain was removed (K_*D*_ of 78 *vs*. 30 μM) (**Figure 4A,B**). The ^15^N,^1^H-HSQC spectrum shows that binding of PY3 induces a stabilization of WW1 in the tandem domain, as demonstrated by a 2-fold increase in visible cross-peaks, several of which representing β-sheet residues (see **Figure 5** discussed below and **Supplementary Figure S4A**). In terms of stability, changes in the CD spectrum of WW1WW2 upon addition of PY3 reveal that the unusually unstable WW1 is subject to further ligand-induced stabilization (**Figure 4D**), in addition to the stabilization by the WW2 in the tandem domain (**Figure 3A)**.

**Table II:**
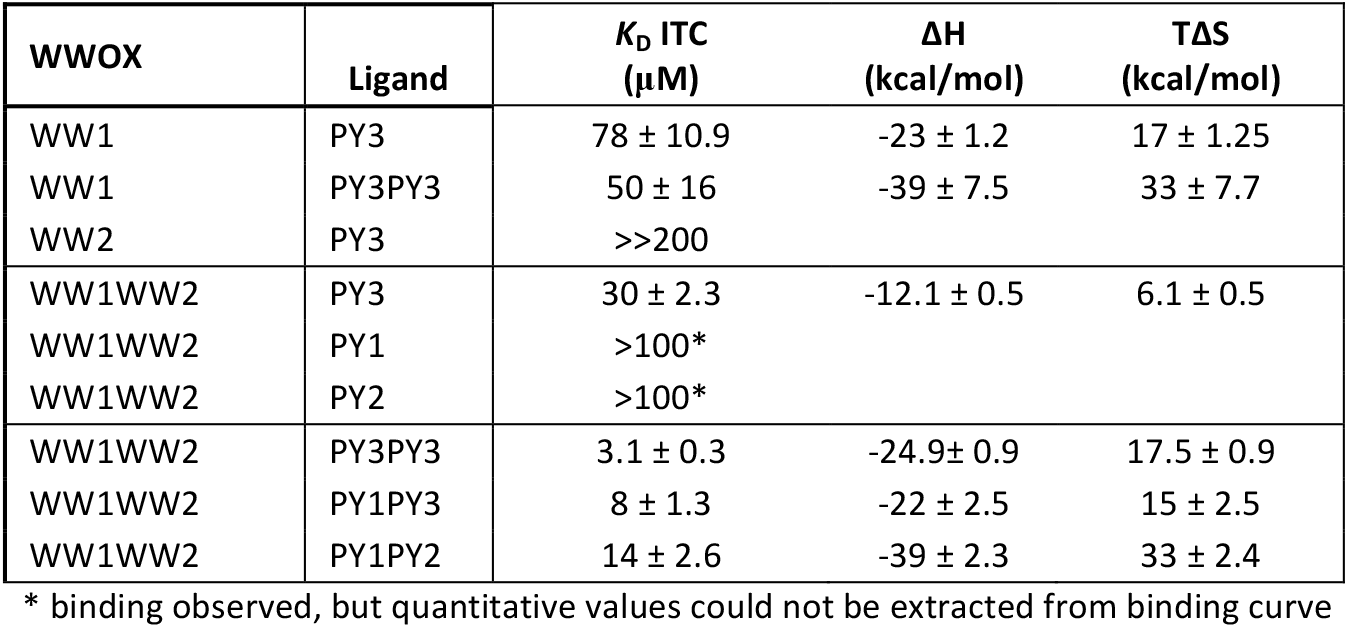
Binding affinities of PPxY peptides to WWOX single and tandem WW domains as determined by ITC (see Experimental Procedures). K_D_ and ΔH/TΔS are given in μM and kcal/mol, respectively. See Table I for details on WW domains and peptide motifs used. Values were compiled from n=3 independent experiments.

| WWOX | Ligand | $K_D$ ITC ($\mu M$) | $\Delta H$ (kcal/mol) | $\Delta S$ (kcal/mol) |
| --- | --- | --- | --- | --- |
| WW1 | PY3 | $78 \pm 10.9$ | $-23 \pm 1.2$ | $17 \pm 1.25$ |
| WW1 | PY3PY3 | $50 \pm 16$ | $-39 \pm 7.5$ | $33 \pm 7.7$ |
| WW2 | PY3 | $>>200$ | | |
| WW1WW2 | PY3 | $30 \pm 2.3$ | $-12.1 \pm 0.5$ | $6.1 \pm 0.5$ |
| WW1WW2 | PY1 | $>100^*$ | | |
| WW1WW2 | PY2 | $>100^*$ | | |
| WW1WW2 | PY3PY3 | $3.1 \pm 0.3$ | $-24.9 \pm 0.9$ | $17.5 \pm 0.9$ |
| WW1WW2 | PY1PY3 | $8 \pm 1.3$ | $-22 \pm 2.5$ | $15 \pm 2.5$ |
| WW1WW2 | PY1PY2 | $14 \pm 2.6$ | $-39 \pm 2.3$ | $33 \pm 2.4$ |

\* binding observed, but quantitative values could not be extracted from binding curve

**Figure 4:**
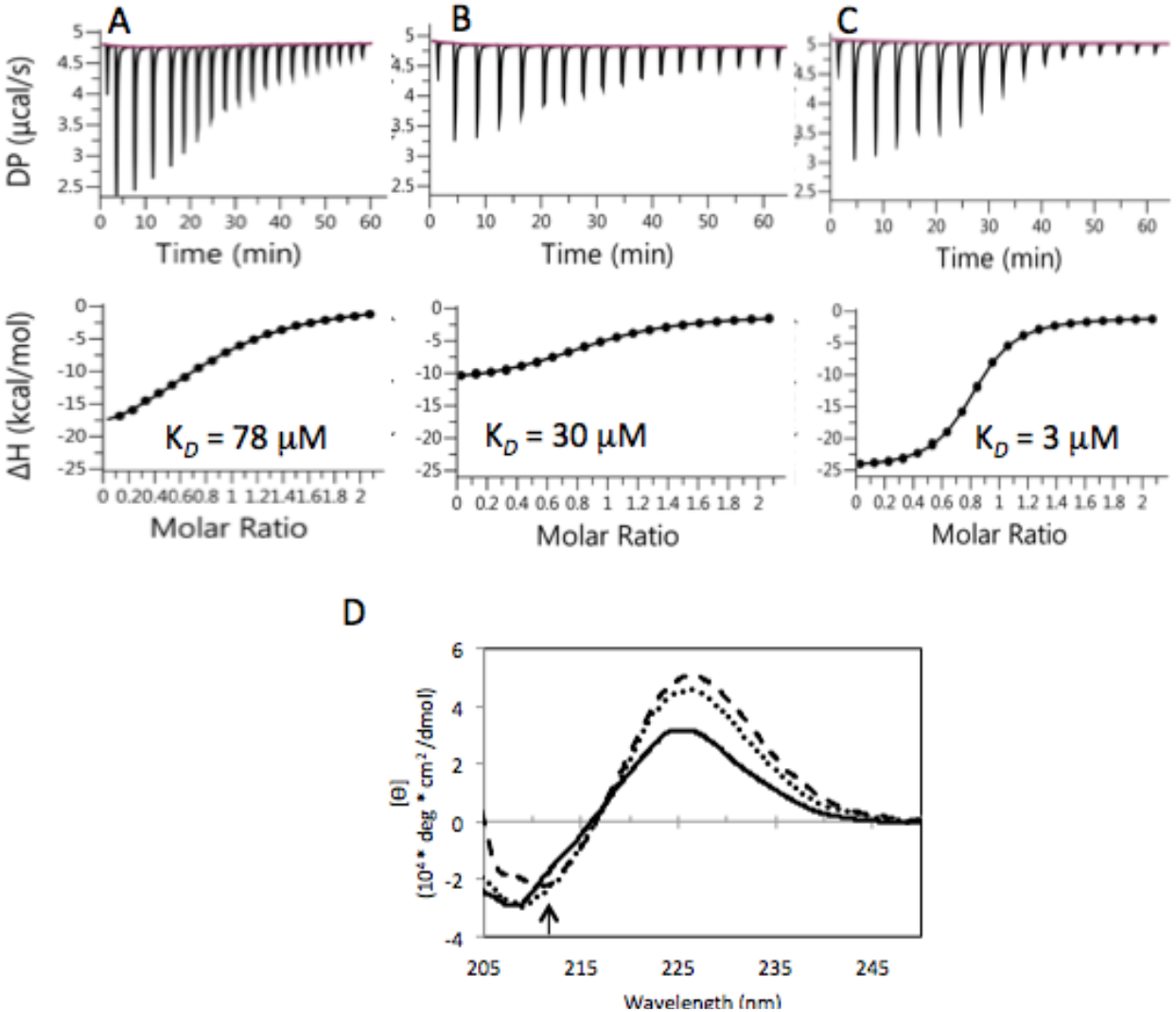
Substrate binding affinity is increased in the presence of WW2 and for a double-PPxY peptide. **A-C**. Isothermal titration calorimetry (ITC) curves of **(A)** WW1, **(B)** WW1-WW2 binding to single motif peptide PY3, and **(C)** of WW1-WW2 binding to double-motif peptide PY3PY3. Calculated affinities are shown. **(D)** CD curves of WWOX WW1-WW2 with no peptide (solid line), when bound to single-motif peptide PY3 (dots), or double motif peptide PY3PY3 (dashed line). The higher intensity at 225 nm is due to increased order in aromatic side chains and to polyproline type II (PPII) helical conformation of the peptide induced by the binding. Peptide CD curves were subtracted to highlight WW contributions to the spectra (For curves before subtraction and curves of isolated peptides, see **Supplementary Figure S2**).

**Figure 5:**
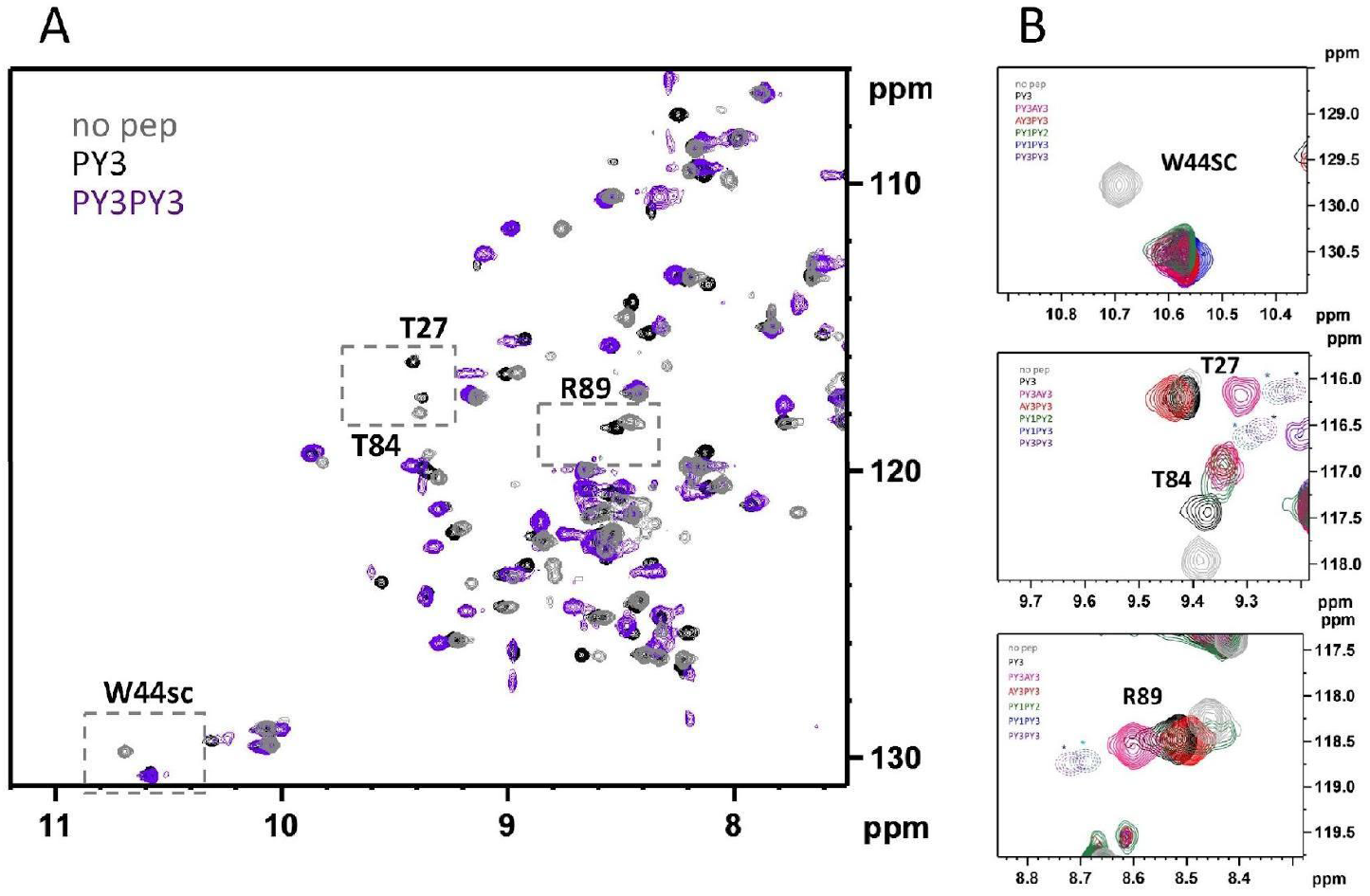
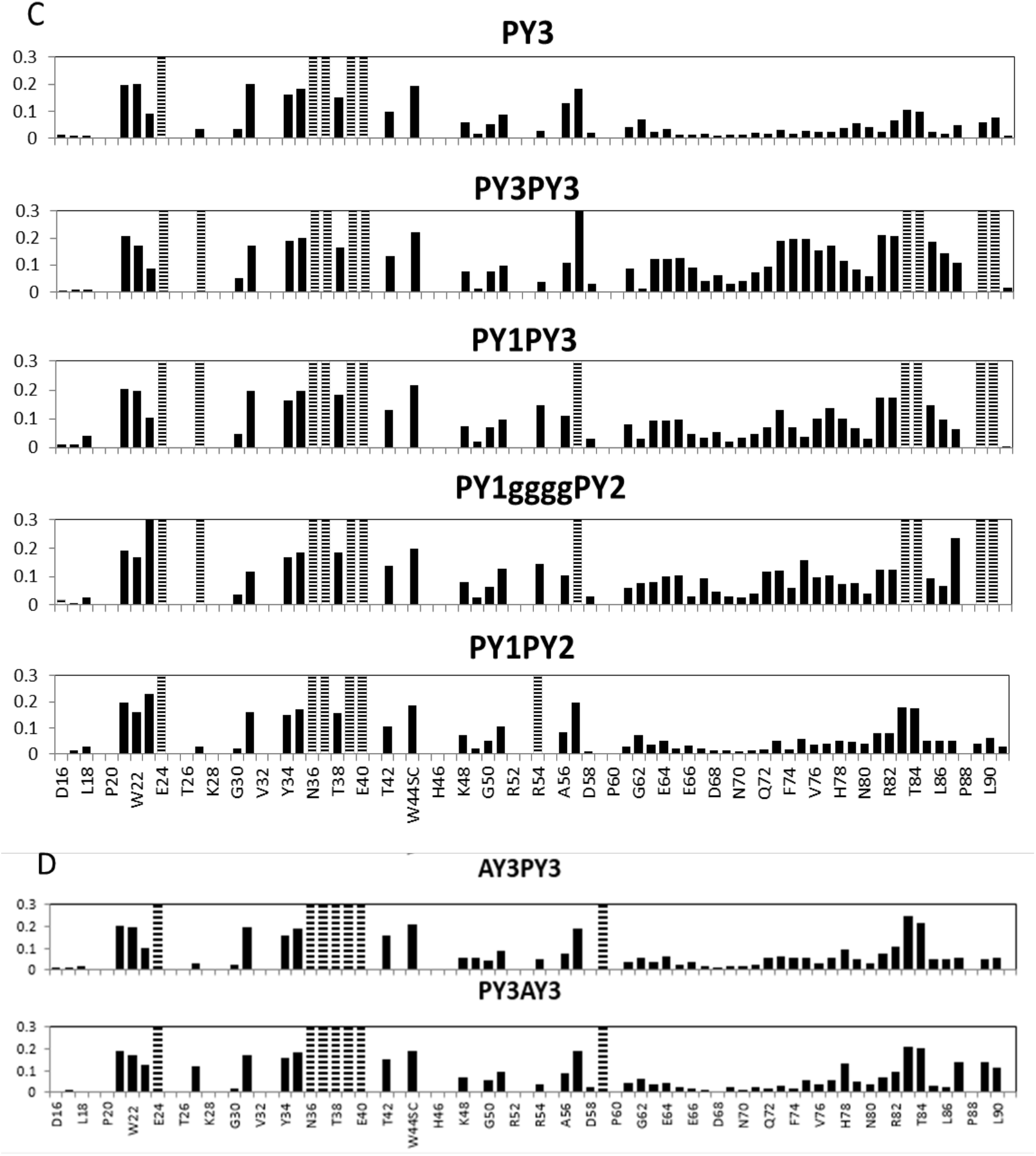
Binding of dual-PPxY peptides to WWOX. **A** Details of ^15^N,^1^H-HSQC spectra of the unbound tandem domains (**gray**) and after addition of single and dual-PPxY peptides at 2:1 peptide:WWOX ratio (PY3 **black**, and PY3PY3 **purple**, respectively). **B** Specific peaks with distinct perturbation patterns upon addition of various peptides, including PY3, PY3PY3, and also PY1PY3 (blue), PY1PY2 (green), PY3AY3 (pink) and AY3PY3 (red) (see Table I for peptide sequences). Peaks that disappear are shown in dotted lines. **C**. Plots summarizing the CSPs observed after addition of 2 molar equivalents of the various peptides. CSP of 0.3 designates a peak that was broadened beyond detection. **D**. Corresponding plots for mutant peptides.

### A significant increase in binding affinity is observed for double binding motifs

While WW domain crosstalk has been extensively discussed (4), less is known about how *binding motif*/multiplicity in the partner affects affinity and specificity. To unravel the effect of motif repeats in the substrate on WW domain binding, we generated several double-motif peptides and measured their binding affinity to WWOX (see **Tables I & II**). In these double-motif ligands the two PPxY motifs are connected by a natural linker segment in the case of the proximal PY1PY2, and a designed poly-glycine linker for the non-native PY1PY3 and PY3PY3 peptides. Such duplication of motifs significantly increased binding affinity, often by an order of magnitude, even when the isolated motifs (*i.e*. PY1 or PY2) failed to bind. Thus, PY1PY2 binds WW1WW2 with a *K*_D_ of 14 μM (as opposed to >100 μM for PY1/PY2 alone), PY1PY3 binds with a *K*_D_ of 8 μM (stronger than PY1/PY3 alone), and PY3PY3 binds with a *K*_D_ of 3 μM (**Figure 4C**, 30 μM for PY3 alone). The PY3PY3 affinity was the strongest measured in this study. This increase in affinity for the dual motif peptides could be seen in CD experiments as well, as these suggest that peptide-induced tightening of the WW domain was more pronounced for double motif ligands (**Figure 4D)**. Although an increase in β-sheet content induced by the peptides could not be sufficiently separated from absorbance of the free peptide molecules (see **Supplementary Figure S2**), these results do suggest that tandem PPxY motif binding induces further structural changes beyond those induced by a single PPxY ligand.

To quantify the contribution of an avidity effect brought about by the two proximal ligands, we measured the binding of PY3PY3 to the single WW1 domain, and observed only a small change in affinity (< twofold). Moreover, a SEC MALS WW1WW2 elution peak had a calculated molecular weight of ca. 10 kDa (**Supplementary Figure S3**), ruling out the possibility that it behaves as a dimer (as occurs, for example, in the SAV1 protein (29)). These results all prove that WW2 plays an active role in peptide binding, and that increased binding affinity is only partially due to avidity.

### Deconvoluting domain contributions to double binding motifs

NMR offers a dual advantage in characterizing the WW/PPxY interactions. First, it is capable of identifying weak affinities (> 100 μM) by following chemical shifts under conditions of exchange which is fast on the NMR timescale. Second, and more importantly, since NMR monitors local changes at each residue, it provides domain-specific binding information, as opposed to the global view afforded by ITC. We exploited this feature of the WWOX NMR fingerprint spectra to analyze the relative contributions of WW1 and WW2 to the binding of double-PPxY peptides. **Figures 5A-B** show the effects of titrating PPxY peptides into a WW1-WW2 sample, and **Figure 5C** summarizes CSPs in the WW1WW2 spectrum for the single motif PY3 and the three double-motif peptides PY1PY2, PY1PY3 and PY3PY3, reporting on contacts between WWOX and the various ligands. The HSQC spectrum (**Figure 5A**) shows single-motif PY3 induced significant peak shifts for most assigned WW1 peaks, in particular for beta strand residues and the W44 indole signal (**Figure 5B**), in agreement with the canonical WW/PPxY binding mode. Concomitantly, only a few WW2 peaks, located in the β_3_-strand, changed significantly upon binding of PY3. Thus, WW1 outcompetes WW2 for PY3 binding, and interaction between the W-deficient WW2 domain and PPxY ligands occurs only due to an increase in local concentration attributable to the canonical WW1/PY interaction. However, changes induced by the three double-motif peptides draw a more complex picture **(Figure 5C)**. The PY3 CSP pattern in WW1 is echoed in the spectra of double-motif peptides, suggesting they all employ a similar binding mode. In contrast, the WW2 CSP pattern distinguished between PY1PY2, reminiscent of PY3, and PY1PY3 and PY3PY3 for which large WW2 CSPs suggest a strong interaction with this domain as well. To some extent this is due to the longer linker connecting PY1 and PY2, since the interaction grew stronger when a shorter flexible non-native poly-glycine linker (similar to the one in the other two double-motif peptides) was used (see **Supplementary Figure S4B**). Overall, the size of the average WW2 CSP was consistent with the affinities of the double motifs, PY1PY2 < PY1PY3 < PY3PY3, suggesting a common WW1 interaction mode and a discriminating WW2 interaction which determines the affinity. For a more quantitative view of domain-specific contributions to affinity, we assumed a two-site binding event with distinct WW1- and WW2-related affinities and followed the concentration-dependent CSP along a titration curve for WW1 and WW2 cross-peaks separately (e.g. the convenient W44 indole and V73 backbone cross-peaks, easily identified in all spectra, see **Supplementary Figure S5**). This resulted in domain-specific values for *apparent* affinities (**Table III** and Experimental Procedures). While PY1PY2 exhibits apparent affinity of ^~^1000 μM to WW2, PY1PY3 and PY3PY3 exhibit appreciable apparent WW2-affinities, 85 and 40 μM, respectively.

**Table III:** Binding affinities of single and dual PPxY peptides to WWOX tandem domain as determined by NMR (see Experimental Procedures). See **Table I** for details on WW domains and peptide motifs used.

| WWOX | Ligand | $K_D$ WW1 ($\mu\text{M}$)<br>Based on W44 | $K_D$ WW2 ($\mu\text{M}$)<br>Based on V73 |
| --- | --- | --- | --- |
| WW1WW2 | PY3 | $10 \pm 2$ | > 1000 |
| WW1WW2 | PY1 | $21 \pm 2$ | ND* |
| WW1WW2 | PY2 | $15 \pm 4$ | ND* |
| WW1WW2 | PY3PY3 | $0.6 \pm 0.3$ | 40 |
| WW1WW2 | PY1PY3 | $6 \pm 1$ | 85 |
| WW1WW2 | PY1PY2 | $10 \pm 2$ | ~ 1000 |

\* ND: Not Detectable

Interestingly, WW2 CSPs induced by binding to the double motif peptides are correlated with additional residues namely WW1 β_1_ residue T27, as well as C-terminal residues R89-L90-A91 (**Figure 5B**), all not located at the peptide binding interface. A structural model of the WWOX tandem domain (see below), co-localizes these four residues at the inter-domain core. It is thus reasonable that these CSPs reflect the bridging of the two domains by the dual motif peptides, a clear indication that two-site binding is occurring. While PY1PY3 and PY3PY3 form two contact surfaces, PY1PY2 binds with a single motif to WW1. Since PY1PY2 does exhibit significant affinity to WW1WW2 despite this difference a more complex mode of binding for this peptide may be indicated.

### Binding orientation of the double motif peptide on the tandem domain

To further elucidate the binding mode of the double motif PY3PY3 peptide to the tandem domain, in particular the directionality of binding (parallel vs. antiparallel), we generated two mutant tandem peptides in which either the N-terminal (termed AY3PY3) or the C-terminal motif (termed PY3AY3) was obviated by PP-to-AA mutations (see **Tables I & IV**; mutation of these residues was shown before to abolish binding (21)). Mutation in the first (AY3PY3) or the second (PY3AY3) peptide motif afforded ITC K_D_ values of 18 μM and 27 μM respectively, closer to the affinity of the single-motif peptide (PY3, 30 μM) than to that of the double motif peptide (PY3PY3, 3μM), indicating non-symmetrical effects of the two mutations on WW1 binding. NMR titration curves afforded poor fits to single-site binding (although K_D_ values in the 10-20 μM range were consistent with ITC results), most likely due to the presence of a residual binding effect of the mutated motif. AY3PY3 did induce slightly larger W44 CSPs (**Figure 5B**), again in agreement with the ITC measurements. On the other hand, of the two peptides PY3AY3 (and not AY3PY3) induced the CSP pattern of inter-domain core residues T27/R89-A91 (**Figure 5D and Figure 6**, top two spectra) previously established as an indicator of domain bridging and two-site binding. A plausible explanation is found in the WW2 CSP values, particularly for residues T83/T84 that are the largest WW2 change observed in most peptides. While these titrations reflect weak WW2/PY interactions in all cases, AY3PY3 curves (but *not* PY3AY3) are bi-phasic (cannot be fitted to a single binding event, **Figure 6**). This can be explained by assuming that single-site WW1/PY3 interactions dominate at low concentrations, whereas at higher concentrations WW1/AY3 interactions increase, allowing some WW2/PY3 encounters to occur. Conversely, for the PY3AY3 peptide the WW1/PY3 interaction is strongest at all concentrations, leaving only a very weak (and monophasic) WW2/AY3 interaction (**Figure 6**). Enthalpy change values observed in ITC measurements are in agreement with this hypothesis, as the PY3AY3 and PY3 share indistinguishable ΔH values (12-13 kcal/mol), whereas the AY3PY3 ΔH (15.6 kcal/mol) indicates an additional interaction (**Table IV**). Using this model to address the question of directionality, we note that PY3AY3, for which the first motif is well-anchored to WW1, exhibits an interaction with the interdomain core, whereas AY3PY3, actually the stronger ligand, draws its higher affinity from a combination of interactions but does not appreciably bridge the two domains. For PY3AY3 this supports a mode in which WW1 interacts strongly with the PY3 motif, and WW2 interacts with the AY3 motif (only at high concentrations due to the weakness of the interaction), and in doing so ‘crosses-over’ the interdomain core, affording a parallel binding orientation. In contrast to AY3PY3, the alternative PY3AY3 binding pose including a WW2/PY3 interaction (as a single domain) appears to be highly unfavorable.

**Table IV:** Binding affinities of peptides mutated in one of the binding motifs. K_D_ and ΔH/TΔS are given in μM and kcal/mol, respectively. See **Table I** for details on WW domains and peptide motifs used. Values were compiled from n=3 independent experiments (except for *: average of 2 experiments).

| WWOX | Ligand | $K_D$ ($\mu\text{M}$) | $\Delta H$ (kcal/mol) | $\Delta S$ (kcal/mol) |
| --- | --- | --- | --- | --- |
| WW1WW2 | PY3 | $30 \pm 2.3$ | $-12.1 \pm 0.5$ | $6.1 \pm 0.5$ |
| WW1WW2 | PY3PY3 | $3.1 \pm 0.3$ | $-24.9 \pm 0.9$ | $17.5 \pm 0.9$ |
| WW1WW2 | AY3PY3 | $18 \pm 3.3$ | $-15 \pm 1.4$ | $9 \pm 1.6$ |
| WW1WW2 | PY3AY3 | $27.1^*$ | $-12.9$ | $6.8$ |

**Figure 6:**
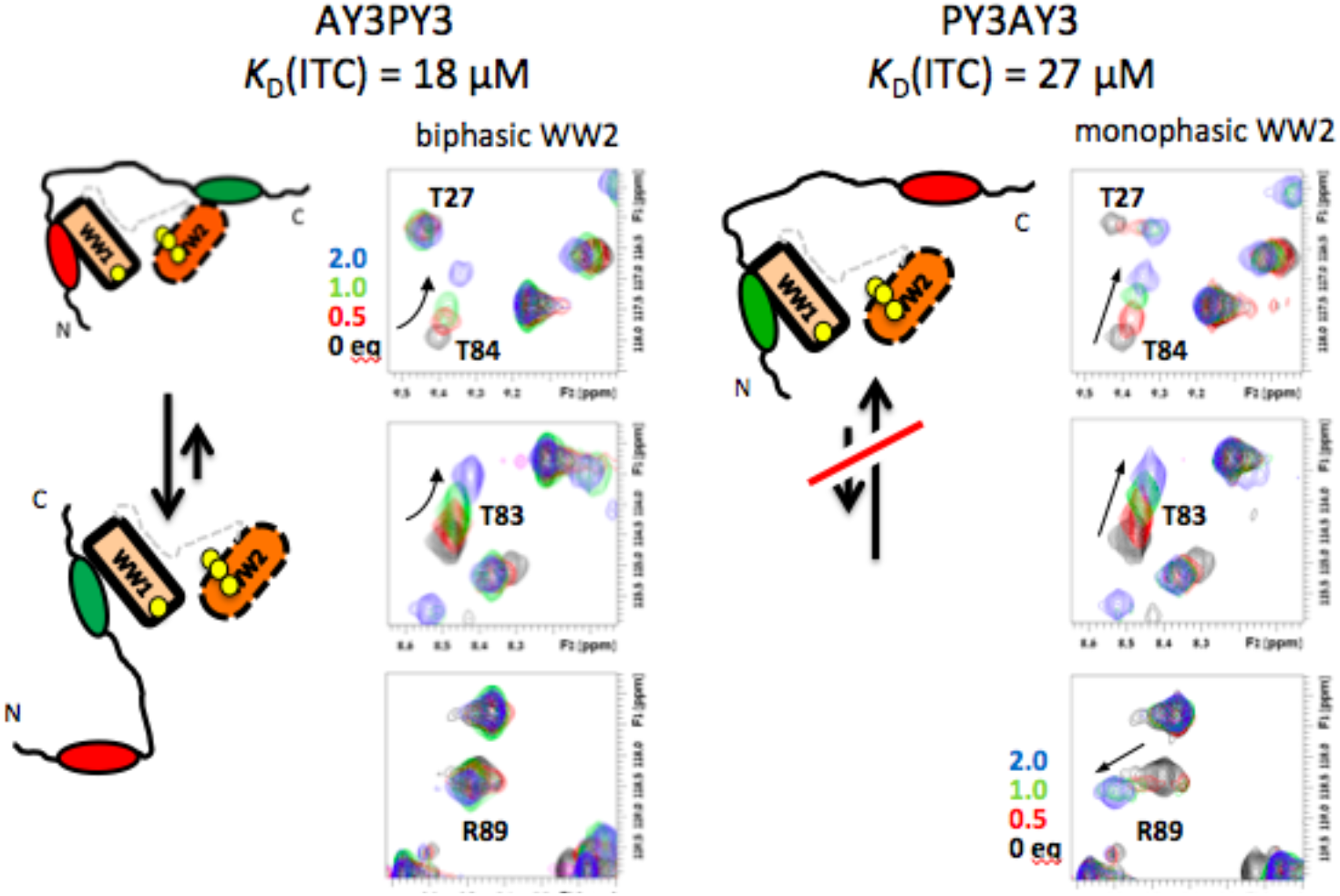
**Scheme of proposed interaction between WWOX tandem WW domain and double motif peptides**, based on distinct effects of mutation of the first or second PPXY motif on binding affinity (from ITC: values at the top; and NMR titration) and perturbation of WW1 and WW2 domains (from NMR HSQC spectra). Models are based on a parallel binding orientation as described in the next section. **Left**, the AY3PY3 peptide with spectral insets showing residues T27, R89, T83 and T84 at 0, 0.5, 1 and 2 mol:mol peptide:WWOX ratios. **Right**, a similar presentation for the PY3AY3 peptide, PY3AY3 shows no sign of a second binding pose (see text). WW1 and WW2 are shown in light green and orange, respectively, The PY3 (AY3) motifs are depicted in dark green (red). Location of residues T27, R89, L90 and A91 is designated with yellow dots. N and C designated the N- and C-termini of the double motif polypeptide.

### Structural models of the WW1WW2 domain

One of the most challenging aspects of understanding the interaction between a double WW domain protein and a double binding motif has been providing a structural view that unifies all experimental findings. While most probably the WWOX WW1WW2 tandem domain adopts an ensemble of interchanging conformations, our experiments still suggest that one or a few will strongly dominate this ensemble. Successful crystallization of such assemblies is plagued by relatively flexible domains (as in the case of WW1) and linkers as well as mediocre affinity levels that hamper such efforts. In light of the combination of the more global ITC/CD/fluorescence experiments alongside NMR experiments affording local per-residue information, we are now in a position to tackle this structural question. In formulating this we draw upon two important conclusions of the current study: (i) Residues T27 (WW1) and R89-L90-A91 (WW2), exhibiting correlated CSPs in all titration experiments, participate in an inter-domain core that stabilizes WW1, and (ii) double-motif peptides interact with both WW domains in a parallel orientation, with the N- and C-terminal PPxY motifs binding WW1 and WW2, respectively.

Thanks to significant recent advances in structure prediction, spearheaded by Alphafold2 (30), and its latest implementation for multimers (31) we were able to generate models that agree surprisingly well with these constraints **(Figure 7A)**. Our model positions the double motif peptide PY3PY3 onto the tandem domain in a parallel orientation, in agreement with our experimental results. Mapping of the electrostatic potential on the surface highlights the acidic patches located near the c-terminus of the PPxY binding motif (**Figure 7B**), explaining the preference in this region for basic residues, as already reported previously (21). In turn a model for PY1PY2 connected by a longer, natural linker positioned only the first motif into the WW1 binding site, while the second motif did not form the canonical interaction between the first proline in the motif and Y85, and moved slightly away.

**Figure 7.**
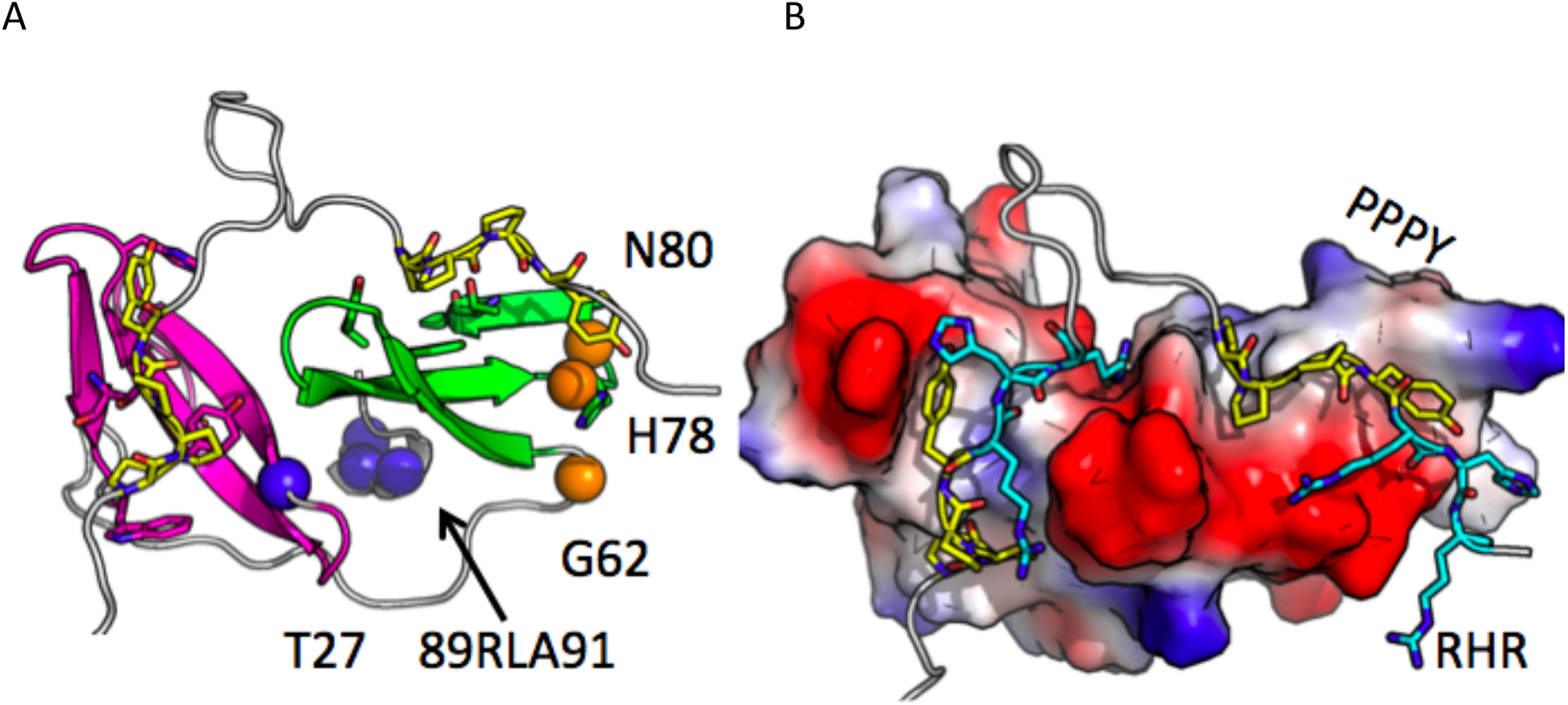
Structural model of the WWOX WW1WW2 tandem domain and its interaction with the tandem PPXY peptide. **A**. Model of WW1WW2 bound to PY3PY3, generated with AlphaFold2. WW1 and WW2 domains are colored in magenta and green, respectively; peptide motifs PPPY are shown in yellow sticks. The parallel orientation adapted by the double peptide motif when binding to the tandem WW domain pair is apparent. The c-terminal residues of WW2 89RLA91 and WW1 T27 are positioned in the center in this model, explaining the concerted shifts observed for these residues upon binding the double motif peptide (see **Figure 5**). In turn, the shifts observed for the beta sheet edge (namely G62 at the beginning of β1, and H78 and N80 in the β2β3 turn), upon addition of WW1 to WW2, could be explained by a change in the beta sheet, or the strain of the linker that is propagated via G62 to that region. **B**. Electrostatic map of WW1WW2, highlighting the strong negative patches, in particular in the WW2 domain, which is contacted by the positively charged flanking regions RHR in the peptide (in cyan). Positions discussed in the text are labeled.

All these features indicate that the tandem domain adopts an open conformation that allows peptide binding, rather than necessitating large rearrangements to free the binding sites. While this suggested model might be the dominant conformation, it is not necessarily the only one (as also indicated by the wide range of different domain-domain orientations that we observe in a larger set of models generated using a number of different approaches, data not shown). The equilibrium between conformations could be shifted to this conformation in particular after binding double motif peptide PY3PY3, while to other conformations upon binding of different peptides, such as PY1PY2.

## Discussion

### WW2 stabilizes the partially unfolded WW1 domain

As many other small peptide-binding domains, WW domains tend to occur in tandem, which allows for the interaction with several corresponding peptide motifs. The ample information on different tandem motif-containing proteins, such as YAP, WWOX and NEDD4 has revealed a wide variety of different strategies that such a framework provides for that purpose (4). WWOX is a particularly interesting example as it takes the way of cooperation to its extreme by optimizing two domains for a different aspect of the interaction, namely stability (WWOX WW2) and binding (WWOX WW1). In this study we have investigated the details of this cooperation using a range of complementary biophysical approaches and modeling. Our first important finding is that while the isolated WW1 domain is unstable, WW2 stabilizes this domain through a network of inter-domain contacts. As a consequence, despite the fact that WW2 lacks significant inherent affinity to PY3, the stabilized WWOX tandem domain binds the ErbB4 PY3 substrate with stronger affinity than an isolated WW1 (78 μM to 30 μM for PY3; a ratio similar to previous reports (21)).

### WW2 participates actively in the binding of dual PPxY motif peptides

Previous studies have reported no detectable binding of isolated WW2 to WWOX-binding peptides derived from different proteins in vitro and in cells (17,18,21,22,25,32). Indeed, we too find that WW2 will not bind a single PPxY ligand independently, and that in tandem it is outcompeted by WW1. However, in our second important finding, using ITC and NMR experiments, capable of detecting intermediate and weaker interactions, we demonstrate that WW2 does participate in the binding of peptides that contain a *dual* PPXY motif. Addition of a second PPXY motif consistently increased affinity by up to tenfold, and this increase cannot be attributed to avidity, since the corresponding affinity increase for the single WW1 domain is less than two-fold (78 μM to 50 μM for PY3 *vs*. PY3PY3) (**Figure 4 & Table II**). This increase in binding affinity necessitates active involvement of the WW2 peptide-binding pocket despite its inherent low affinity for peptide substrates (due to the missing canonical tryptophan residue), presumably due to increased local effective concentration (**Figures 4–6**). It is true that such effects are seen in other systems, e.g. tandem SH2 domains bind double phospho-tyrosine peptides with >1000-fold higher affinity than a single domain (33), and the fibronectin (Fn) N-terminal domain contains five type 1 modules and binds short repeat motifs in the SfbI protein of *Streptococcus pyogenes* (34)). However, in most of these and other cases, both domains bind the PPxY motif in isolation. Here we find a dual role for WW2: A stabilizing influence on the partially unfolded WW1 domain as well as substrate binding in the case of double-PPxY motifs.

### A molecular view of the parallel tandem-WW/dual-PPxY assembly

The essence of biological regulation by the interactions of WW domains and their PPxY ligands is the ability of a tandem WW domain to capture polypeptides containing proximal PPxY sequences. We explored the molecular factors governing this binding interaction for WWOX and a series of designed polypeptides. As expected, one factor is the inherent affinity of the PPxY sequence, adhering to previously reported relative binding strengths (PY3 > PY1 > PY2, determined by the presence of a key basic residue following the PPxY motif (21)). However, a second factor is the relative location of the two binding sequences. The connecting linker length is a well-recognized determinant of tandem binding affinities, as in polypeptides with multiple PPxY motifs binding predominantly occurs for proximal motifs (separated by ^~^15-20 residues), and only rarely for distal motifs (5,35). These two factors embody enthalpic and entropic contributions, respectively, to binding. Here we report a significant difference in the binding mode of the PY3PY3 and PY1PY3 peptides compared to PY1PY2, where WW2 is bound appreciably only by the former, while the latter binds predominantly to WW1 (**Figure 5**). Replacement of the native linker by a short polyglycine sequence in PY1PY2 (as in PY1PY3 and PY3PY3) did lead to stronger interaction with WW2 (**Figures 5C & S4B**).

Our third important finding is the establishment of a parallel binding mode, by investigating the effect of independently ‘knocking-out’ each of the PPxY motifs in the PY3PY3 peptide. The first PPxY interacts with WW1 and the second binds weakly to WW2 (**Figure 6**). This binding mode is supported by a structural model of the WWOX tandem WW domain interacting with the PY3PY3 double motif peptide (**Figure 7**). Our model reconciles our experimental findings: (1) It reconfirms parallel binding of the peptide to WWOX, positioning the first motif into the first WW1 domain, (2) It provides an explanation for binding to WW2, where a strong electrostatic acidic patch attracts the basic region that flanks the binding motif. Thus, even without a canonical tryptophan, this domain can bind to a peptide motif brought into proximity by its adjacent second motif bound to WW1. (3) It is consistent with prearrangement in a peptide-binding compatible orientation. This conformation is able to bind the double motif peptide in parallel orientation, where the two peptide motifs wrap around WWOX, bringing the two domains closer. Thus, residue T27 in WW1 at the domain interface, as well as the c-terminal region of WW2 are only perturbed by peptides binding also to WW2 (**Figure 6**).

Of note, our model differs from a model suggested in a previous study, which was generated using molecular dynamics simulations starting from the template structure of FBP21 (25,36). That model suggested that the WW1 peptide-binding site is occupied by the WW2 domain and consequently, binding of peptide substrates would involve major domain rearrangement to allow access of the peptide to the binding site. Our NMR results do not provide any support for such a rearrangement, and moreover, regions that show significant change in surrounding when comparing the single and tandem WW domains (as reflected by our measured chemical shift perturbations) are not located at the domain interface.

Beyond the main model presented in **Figure 7** however, WWOX is functional as a dynamic ensemble of different conformations, as suggested from our NMR experiments, as well from previous studies (25,36). While the models generated by AlphaFold2 converge, models that we generated using additional protocols (including Rosetta *ab initio* folding with GREMLIN constraints (37), TrRosetta (38) and RoseTTAFold (39), data not shown) exhibit a range of additional possible relative orientations between WW1 and WW2, with predominantly similar conformation of the individual WW domains, but a wide variability for the linker and the tails. While only few comply with our experimental results, as an ensemble they may set the stage for a more detailed investigation of the possible contribution of different conformations to distinct functional contexts. After all, even if some specific conformations dominate, it is often the ensemble of conformations of a protein that will define its function (40).

### Evolution of tandem domain binding: WWOX adopts a new WW binding mode

This study and others have closely examined the structural factors governing binding affinities between tandem WW domains and their single (or dual) PPxY ligands. Comparison of tandem WW domains demonstrates that their interaction with their natural ligands is controlled by a multifactorial array of sequence- and structure-related parameters. This is schematically summarized in **Figure 8A**. Variants of the WW domain - stable, ‘unfolded’, and mutated (e.g. loss of tryptophan) - exhibiting different inherent PPxY affinities, are connected by linkers differing in length and structural flexibility. We find WWOX to be relatively unstructured in WW1 (similarly to Su(dx)), lacking the canonical tryptophan in WW2 (as in Su(dx) and KIBRA), and connected by a flexible linker (as in FBP21, shorter than the YAP linker, **Figure 8B**). A degree of coupling between the two domains is observed, with significant WW2-induced stabilization of WW1, although effects of PPxY binding to WW1 are not propagated to WW2. Despite these partial similarities, binding of dual PPxY by WWOX does not fully resemble any solved structure of a tandem WW domain - peptide interaction (**Figure 8C**).

**Figure 8:**
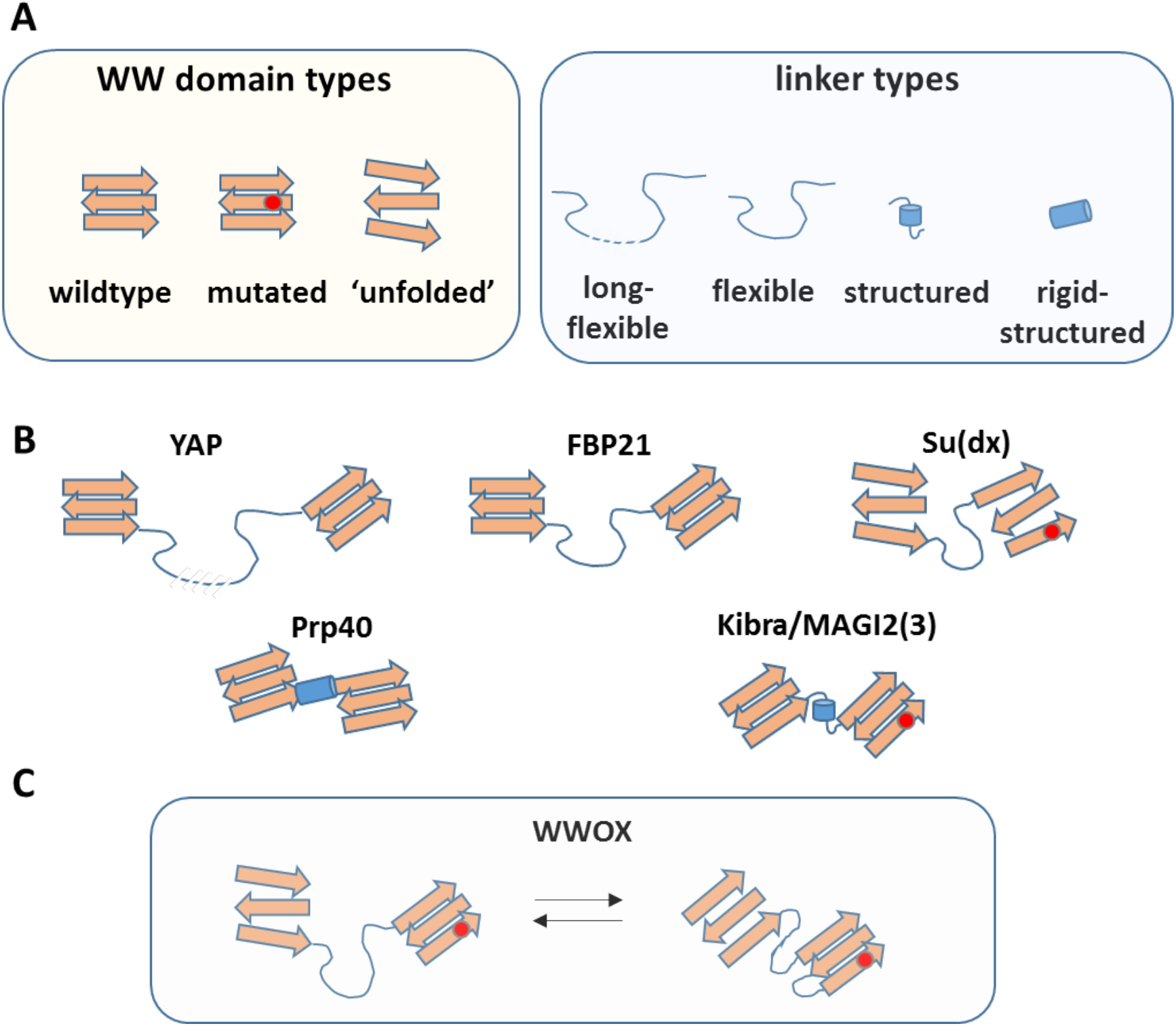
Multi-layer modulation of WW-ligand interactions and affinities. **A**. Components of modulation including the nature of the WW domain (wildtype, mutant, ‘unfolded’) and the linker domain (length, flexible/rigid). **B**. Schematic representation of known tandem-WW structures (not necessarily an exhaustive list) using the above building blocks. Shown are YAP (41), FBP21 (7), Su(dx) (42), Prp splicing factor (43), and KIBRA (which is similar to protein MAGI2(3)) (10,41). **C**. Representation of WWOX, the subject of this study, using the same building blocks. The presence of WW2 exerts a stabilizing effect on WW1 via interactions with the linker. Similar stabilizing effects occur for other proteins (not shown for clarity).

Additional modes may be considered as well. Just as the Tondu domain-containing growth inhibitor (tgi) protein bears one PPxY motif far from two “low affinity” close motifs (5), the PY3 of ErbB4 may be the high affinity “binding initiation” site leading to interaction to the “low affinity” PY1/PY2 site. There are cases where tandem motifs binding does not improve binding, as for example in the YAP-LATS interaction, where the tandem motif binds with affinity similar to that of the single motif (44). This suggests that this kind of binding interplay does not always play a role in the regulation. Finally, post-translational modifications may also modulate affinity (19,45). The result is a range of seemingly similar tandem interaction modules actually capable of a wide range of affinities, and, commensurately, biological functions. Together with factors present in the ligand, such as variations in the binding motif and the inter-motif distance, all of these impact the balance of enthalpic and entropic factors that eventually determine the nature of the ensuing interaction between tandem WW domain and PPxY ligands.

We find this consistent from an evolutionary viewpoint – since WW-PPxY interactions (barring the missing tryptophan residue in some domains, and occasionally leucines replacing the first proline of the PPxY motif) are relatively similar in structural determinants and consequently in affinity, other factors have evolved to allow a fine-tuning of these interactions in a setting-dependent manner. Here WWOX provides a case in point, exemplifying that Nature can create a less stable, unstructured domain for modifying binding affinity (and possibly specificity) and compensate for this destabilization by an adjacent chaperone domain. This is in agreement with a large-scale analysis of the (calculated) stability of single domains *vs*. their corresponding occurrence within a multi-domain setting. Domains that lack independent stability were stabilized by favorable interactions with other domains, stabilization of such folds was optimized by evolution, and specific mutations at inter-domain hotspots could rescue the overall stability (46), again underlining the importance of non-canonical factors as an evolutionary tool for fine-tuned control of this family of protein-protein interactions.

To summarize, we have mapped in this study the mutual influence of WW1 and WW2 tandem domains, and their involvement in the binding of single and double motif containing peptide substrates in WWOX, using a number of complementary experiments including ITC binding experiments, NMR CSP changes and titrations, structural modeling and more. We have shown a dramatic increase in binding affinity of double motif containing peptide substrates, and demonstrated that WW2 plays an active and important role in the binding of the second peptide motif, beyond merely increasing WW1 stability. These results join and confirm others showing that WW domains have evolved to utilize several structural and sequence factors to modify the affinity for their canonical ligands. Further studies are needed to determine how this combination of two WW domains, one with reduced binding and one with reduced stability, plays an important role in defining the binding specificity, and consequently functionality of WWOX within cells.

## Experimental Procedures

### Expression Plasmids

PCR products WWOX ww1-ww2 (amino acid 16-91), WWOX ww1 (amino acid 16-50) and WWOX ww2 (amino acid 57-91) were cloned into pETM 30 YAP 171-264 plasmid (a kind gift of Dr. Maria J. Macias, Institute for Research in Biomedicine, Barcelona) containing an N-terminal HisX6 tag followed by a GST and TEV protease, instead of the YAP sequence in the NcoI and HindIII restriction sites.

### Protein Expression and Purification

all the WWOX constructs were expressed in *Escherichia coli* BL21 pLysS cells (Novagen). Cells were grown in 2× YT medium. Induction was done at A600 nm = 0.6-0.8 with 0.1 mM isopropyl β-D-thiogalactopyranoside (IPTG) and cells were grown overnight at 20°C. Isotopically labeled proteins for NMR measurements were expressed in M9 minimal medium (47) supplemented with 1 g/l ^15^NH_4_Cl for ^15^N. ^15^N,^13^C-double labeled samples were also grown with 2.5 g/l ^13^C6-D-glucose (only WWOX ww1-ww2) to OD600=0.8, induced with 0.15 mM IPTG and grown overnight at 27 °C. Cells were harvested by centrifugation and stored at −80 °C.

Cell pellets containing expressed WWOX constructs were resuspended in lysis buffer (50 mM NaH_2_PO_4_ pH 7.0, 300 mM NaCl, 10 mM imidazole, and 5 mM β-mercaptoethanol), supplemented with 1 mM phenyl-methyl sulphonyl fluoride (PMSF) and DNase. Cells were disrupted using a microfluidizer (Microfluidics). Lysate was cleared by centrifugation and was subjected to 5 ml His-Trap columns (GE Healthcare). The protein was eluted with a linear imidazole gradient of 15–250 mM in 30 column volumes. Fractions containing the purified protein were pooled and dialyzed overnight at 4 °C against dialysis buffer (20 mM NaH_2_PO_4_ pH 7.0, 150 mM NaCl, and 5mM β-mercaptoethanol) in the presence of TEV protease. Cleaved protein was then subjected to a second round of His-Trap column and flow-through containing the cleaved protein was collected. The proteins were further purified using 16/600 Superdex 75 pg size-exclusion chromatography columns (GE Healthcare) equilibrated in protein buffer (20mM Tris pH 7.8, 150mM NaCl). NMR samples were prepared in 20 mM NaH_2_PO_4_ buffer, pH 6.8, 100 mM NaCl, 7% ^2^H_2_O and supplemented with EDTA-free protease inhibitor cocktail (Roche). Final protein concentrations were 0.3-0.6 mM. All proteins were concentrated, flash-frozen in liquid nitrogen, and stored at −80 °C.

### Peptide synthesis

peptides were purchased from PHTD peptide (Hong-Kong, China) and GenScript (HK limited, Hong-Kong, China) with 90%-95% purity.

### CD

Circular dichroism spectra of 50 μM WWOX 16-91, WWOX 16-50 and WWOX 57-91 were recorded using a J-810 spectropolarimeter (Jasco) in protein buffer, in a quartz cuvette for far-UV CD spectroscopy. Far-UV CD spectra were collected in a spectral range of 190 to 260 nm. For measuring the spectra of interactions, 50 μM WWOX 16-91 was incubated with 50 μM ErbB4 1291—1305 (PY3) or with 50 μM ErbB4 PY3PY3 in protein buffer. Background scans of the peptides were conducted in buffer alone, and subtracted from the protein with peptide scans.

We note that our CD spectra of WWOX WW domain fragments differ from those reported previously by others (22). This is mainly due to the use of a slightly different construct (including only four extra residues at its N terminus), and the performance of the experiments at different buffers and temperatures.

### Protein Fluorescence

5 μM protein in protein buffer was incubated with or without 6 M urea. Protein tryptophans were excited at 295 nm and emission was measured at 325-400 nm in 96 well plates with Cytation3 imaging reader (BioTek).

### ITC

Isothermal titration calorimetry measurements were performed on an ITC200 calorimeter (Microcal, GE Healthcare) at 20°C. 1-2 mM PY peptides were titrated to 100-200 μM of the different WWOX constructs in protein buffer. The data were fitted using ORIGIN 7.0 software (Origin Lab) (and microcal PEAQ-ITC) to the single-site binding isotherm and N was set to 1 when the K_*D*_ was weaker than 50 μM. The integrated peak of the first injection was excluded from the fit due to the large errors in the first step. Each experiment was repeated at least three times. We note that previous studies have reported slightly different absolute binding affinities (21), mostly due to differences in buffer conditions, but the relative affinities are unchanged.

### SEC-MALS

Size Exclusion Chromatography - Multi Angle Light Scattering Experiments were performed with a pre-equilibrated analytical SEC column (Superdex 200 10/300 GL; GE Healthcare Life Sciences) with protein buffer, as described in Mashahreh *et al*. (48).

### NMR

Nuclear magnetic resonance spectra were recorded on a DRX700 Bruker spectrometer using a cryogenic triple-resonance TCI probehead equipped with z-axis pulsed field gradients. Spectra were measured at 16.4 T and 286 K. ^1^H,^15^N-HSQC spectra for sample characterization and optimization were run for 30-40 minutes with acquisition times of 91.4 (70) ms and 1024 (100) complex points in the F2 (F1) dimensions, respectively. Triple-resonance HNCO, HNCA, HN(CO)CACB, and HNCACB spectra, using sensitivity-enhanced echo–antiecho detection, were acquired for uniformly ^13^C,^15^N-labeled samples. All triple-resonance and ^15^N-edited experiments were typically acquired with 40-48 complex points and an acquisition time of 20-24.1 ms in the ^15^N dimension, and with 1024 complex points and an acquisition time of 91.8-104.4 (145) ms in the observed proton (^13^C’) dimension. For indirect ^13^C dimensions, experiments with ^13^CO(^13^Cα) evolution were acquired with 60 (44) complex points and an acquisition time of 28.3 (7.8) ms, and experiments with ^13^Cα/β evolution were acquired with 44–52 complex points and an acquisition time of 4.1-4.8 ms. Peak assignment was based on these triple resonance spectra; in cases of ambiguity and/or significant peak broadening the assignment was assisted by data acquired for a WW1 A35V mutant that exhibited reduced exchange broadening (data not shown). Processing and analysis of all spectra was performed using the TopSpin 3.2 package (Bruker BioSpin, Karlsruhe, Germany).

Binding of ErbB4 peptides to the proteins was monitored by repeating the ^1^H,^15^N-HSQC spectrum after serial additions of the desired peptide up to 2-4 molar equivalents taking care to maintain a constant WW-domain concentration. Using these spectra the affinity could be estimated by plotting the chemical shift perturbation as a function of the number of peptide equivalents added (49)(see **Supplementary Figure S4**).

### Generation of structural models of WWOX

Structural models of the tandem domain were generated with DeepMind Alphafold2 (31), using the deepmind colab setup (https://colab.research.google.com/github/deepmind/alphafold/blob/main/notebooks/AlphaFold.ipynb). The peptide was provided as a separate chain. We also generated additional models using a range of different approaches, including RoseTTAFold (http://new.robetta.org)(39), TrRosetta (https://yanglab.nankai.edu.cn/trRosetta) (38), and ab initio Rosetta folding under constraints derived by GREMLIN (http://gremlin.bakerlab.org) (37). None of these models provided good agreement with all experimental results (data not shown).

## Data Availability

Raw data files from experiments, as well as structural models, that form the basis of the results presented in this study are provided upon request by the corresponding authors.

## Supporting information

This article contains supporting information: five supplementary Figures (no additional references).

## Acknowledgments

We thank Amjad Farooq and Rami Aqeilan for providing the WWOX plasmids.

## Funding and additional information

This work was supported, in whole or in part, by the Israel Science Foundation, founded by the Israel Academy of Science and Humanities (grant number 717/2017 to O.S.-F., grant number 964/19 to J.H.C.), and from the European Research Council under the ERC Grant Agreement [310873].

## Supplementary Figures

**Figure S1:**
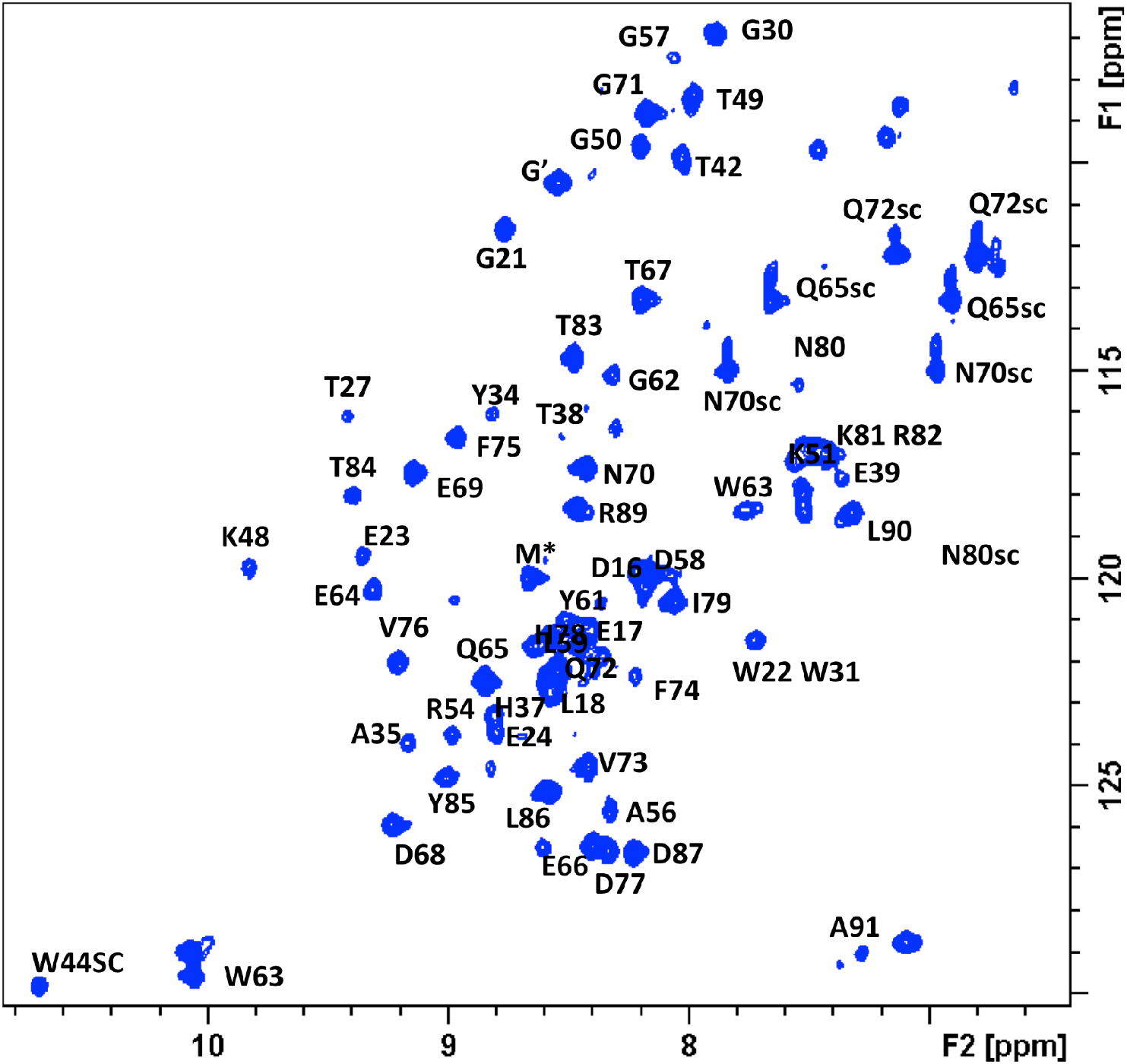
Peak assignments. Shown is the annotated spectrum for WWOX tandem WW1-WW2 domains spanning residues 16-91 (**see Table I**).

**Figure S2:**
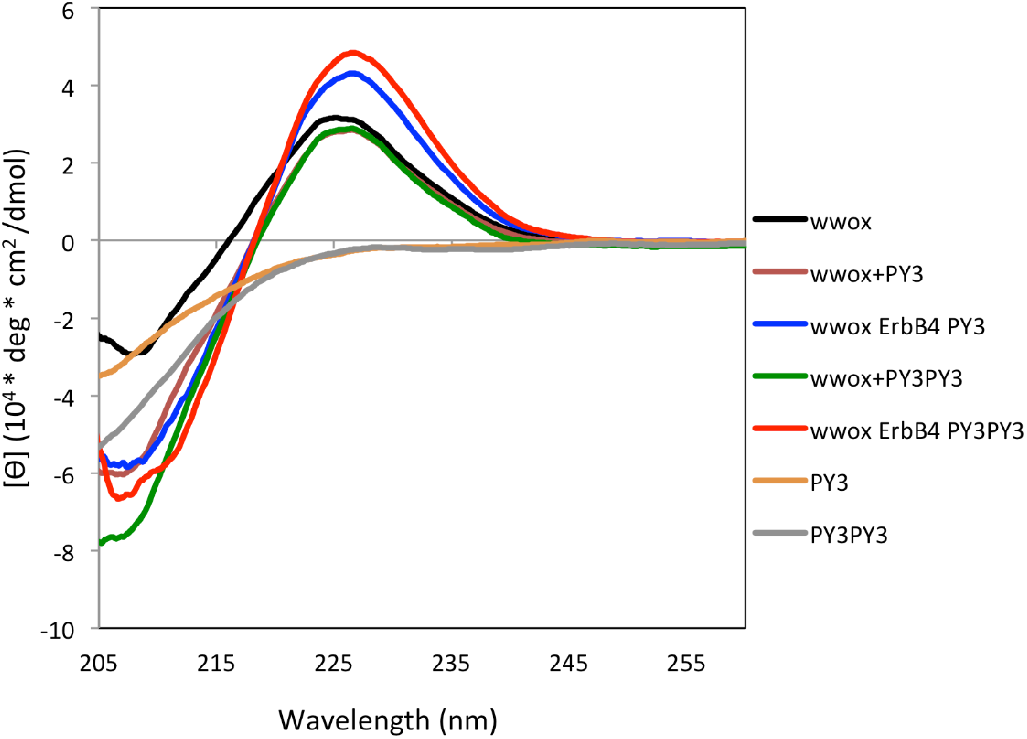
CD analysis of free and peptide-bound WWOX. (accompanies Figure 4D). Shown are the curves (prior to subtraction of peptide contribution) of free WWOX (black, as in **Figure 6D**) in comparison to curves of free single- (brown) and double-motif (green) peptides alone, and curves for the WWOX complex with the peptides (blue and red, respectively). The significant difference of the single and double motif peptide spectra in isolated form in the far left range indicate that some of the observed changes in the spectrum after subtraction are also due to conformational changes in the peptide.

**Figure S3:**
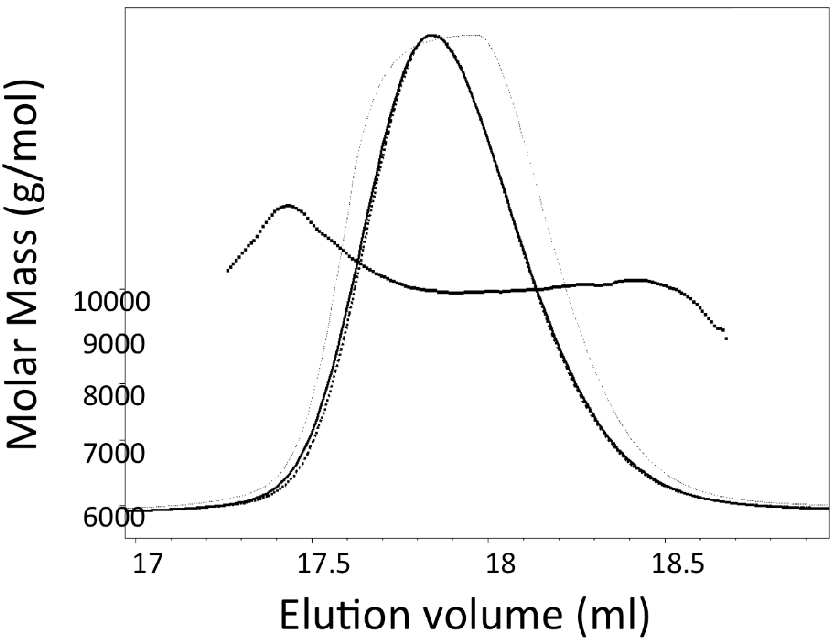
SEC-MALS analysis of WWOX. SEC-MALS shows that WWOX (*i.e*., tandem WW1-WW2) is monomeric with a molecular weight of ~10 kDa as expected.

**Figure S4:**
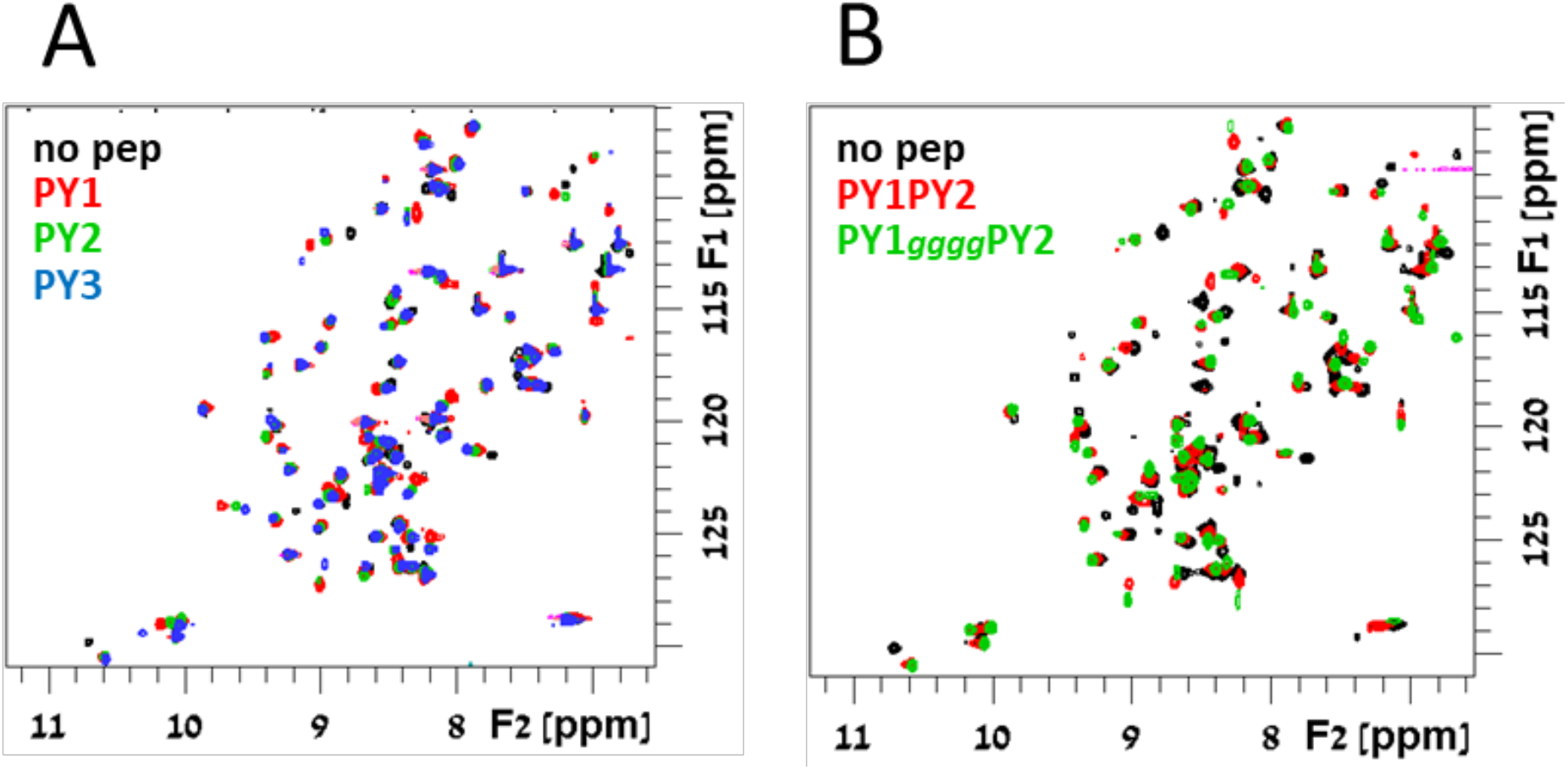
Binding of PY1 and PY2 to WWOX as single motifs and in tandem (accompanies Figure 5). **A**. Details of ^15^N,^1^H-HSQC spectra of the unbound tandem domains (**black**) and after addition of single motif peptides at 2:1 peptide:WWOX ratio (PY1 **red**, PY2 **green**, and PY3 **blue**). **B**. Same as **A** for dual PPxY peptides (native PY1PY2 **red**, and PY1*gggg*PY2 with polyglycine linker **green**, respectively).

**Figure S5:**
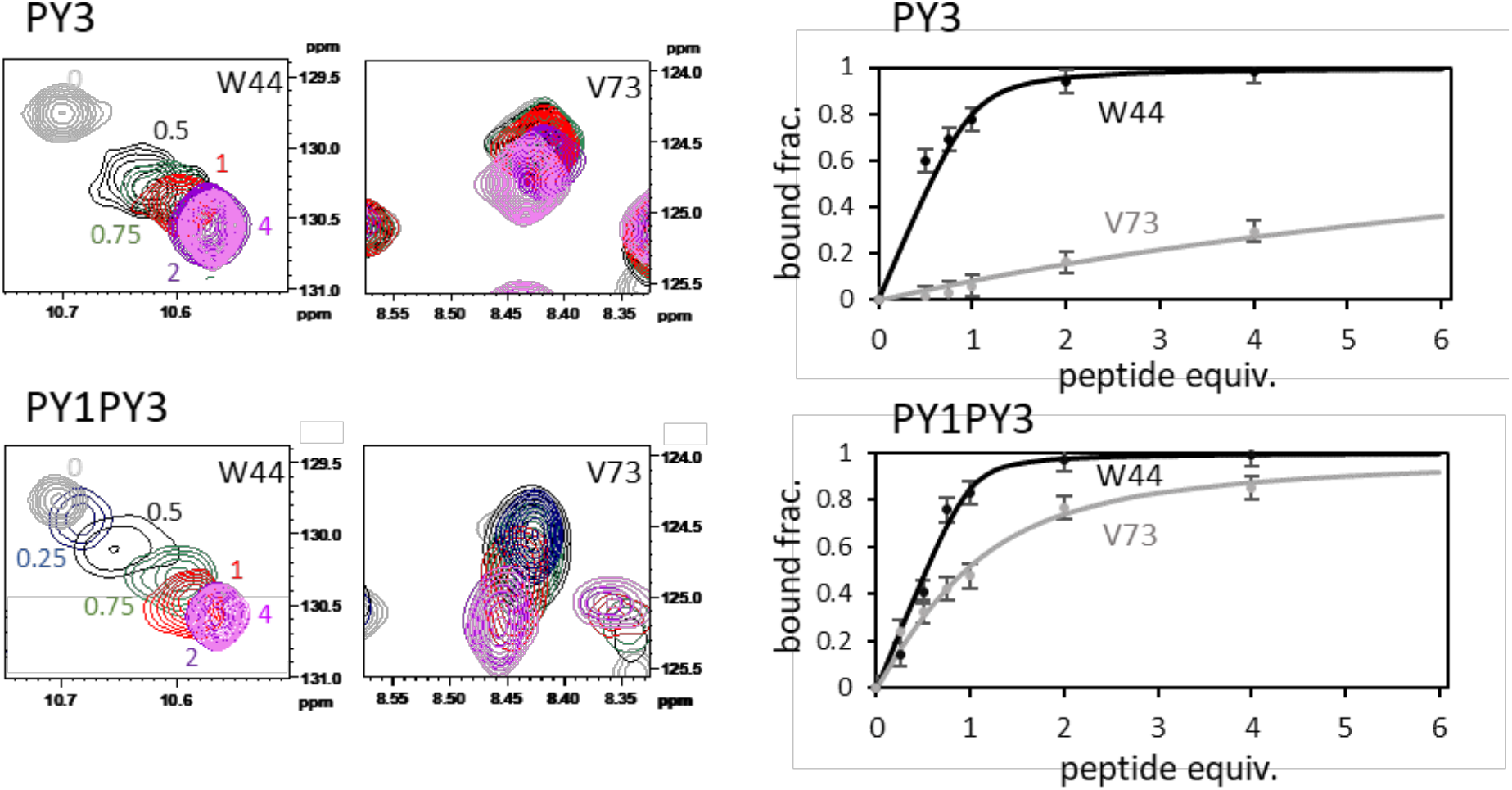
NMR titration experiment for the determination of *K*_D_ values for binding the individual WW domains. Shown are CSP changes upon addition of increasing concentrations of PY3 (top) and PY1PY3 double motif (bottom), for peaks associated with residues W44 (side chain) in WW1 (left), and V73 in WW2 (right). Far right shows the fraction of bound WW1 (WW2) as a function of the number of peptide equivalents added in black (grey) by following changes in W44 (V73). Points are measured data, lines represent best fits to the isotherm equation (see ref. 47).

